# Seasonal influenza circulation patterns and projections for Feb 2018 to Feb 2019

**DOI:** 10.1101/271114

**Authors:** Trevor Bedford, Richard A. Neher

## Abstract

This report details current seasonal influenza circulation patterns as of Feb 2018 and makes projections up to Feb 2019 to coincide with selection of the 2018-2019 Northern Hemisphere vaccine strain. This is not meant as a comprehensive report, but is instead intended as particular observations that we’ve made that may be of relevance. Please also note that observed patterns reflect the GISAID database and may not be entirely representative of underlying dynamics. All analyses are based on the nextflu pipeline [1] with continual updates posted to nextflu.org.

**A/H3N2:** H3N2 diversity has largely been replaced by subclades A1b, A2 and A3 within 3c2.A. Subclades A1b and A2 predominate in the population and each shows increases in frequency, mutations at epitope sites and evidence for minor changes to antigenic phenotype. Clade A1b may be marginally fitter than clade A2, but we expect both clades to persist into the future without a clear immediate winner.

**A/H1N1pdm:** A clade comprising mutations S74R, S164T and I295V has recently swept to fixation. The rapidity of this sweep suggests a selective origin. However, there is no evidence of antigenic change.

**B/Vic:** Very little B/Vic activity has been observed in recent months. A clade with a two codon deletion at sites HA1:162/163 has gradually risen in frequency. HI measurements suggest an 8 to 16-fold titer drop relative to the vaccine strain, but this antigenic change has not yet resulted in a rapid rise of this variant.

**B/Yam:** Europe experienced a strong and early B/Yam season in absence of amino acid variation in HA or antigenic diversity. However, several mutations in NA have rapidly swept or risen to intermediate frequencies.

## A/H3N2

H3N2 diversity has largely been replaced by subclades A1b, A2 and A3 within 3c2.A. Subclades A1b and A2 predominate in the population and each shows increases in frequency, mutations at epitope sites and evidence for minor changes to antigenic phenotype. Clade A1b may be marginally fitter than clade A2, but we expect both clades to persist into the future without a clear immediate winner.

We base our primary analysis on a set of viruses collected between Mar 2016 and Jan 2018, comprising >1000 viruses per month in Dec 2016 to Mar 2017 and 100-200 viruses per month more recently (Fig. 1). We use all available data when estimating frequencies of mutations and weight samples appropriately by regional population size and relative sampling intensity to arrive at a putatively unbiased global frequency estimate. Phylogenetic analyses are based on a representative sample of about 2000 viruses.

**Figure 1.**
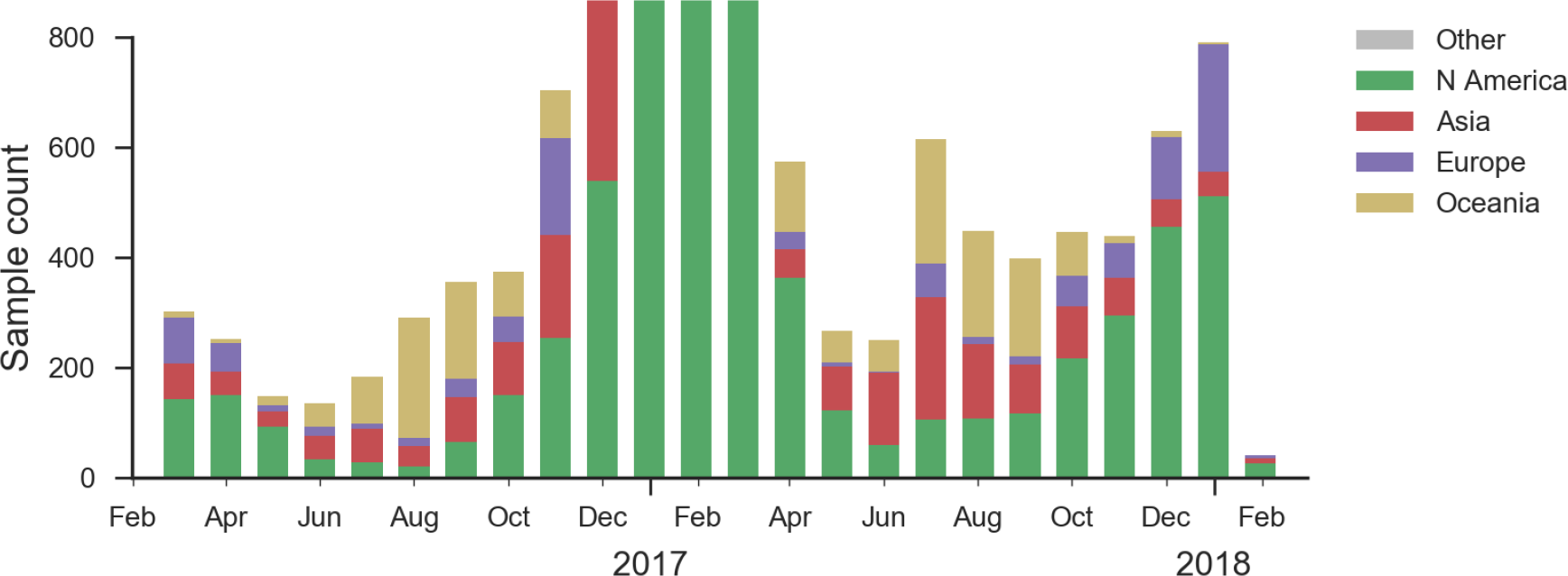
Sample counts through time and across regions. This is a stacked bar plot, so that in all months there are at least 100 samples and in some months there are >1000 sequences.

## Current circulation patterns

Clades 3c3.A and 3c3.B have hardly been observed for over 1 year and the great majority of H3N2 viruses now fall into clade 3c2.A or its subclades. Within 3c2.A, the clade 3c2.A1 (bearing mutation N171K) is now globally at a frequency of about 40%, while North America is dominated by viruses outside this subclade.

We observe substantial diversification within the 3c2.A clade over the past year. There are 5 major subclades within 3c2.A that have reached appreciable frequency (Fig. 2), three deriving from basal clade 3c2.A and two deriving from clade 3c2.A1. All of these clades possess mutations at previously characterized epitope sites. These clades are characterized by the following mutations and highlighted in the phylogenetic tree in Fig. 2:

1. Clade 3c2.A1a: 135K, HA2:150E (previously called clade 4)
2. Clade 3c2.A1b: 92R, 311Q (previously called clade 5)
3. Clade 3c2.A2: 131K, 142K, 261Q (previously called clade 3)
4. Clade 3c2.A3: 121K, 144K (previously called clade 2)
5. Clade 3c2.A4: 53N, 144R, 171K, 192T, 197H (previously called clade 1)

Throughout the following, we drop the 3c2 to abbreviate these names to A1a, A1b, A2, A3 and A4.

**Figure 2.**
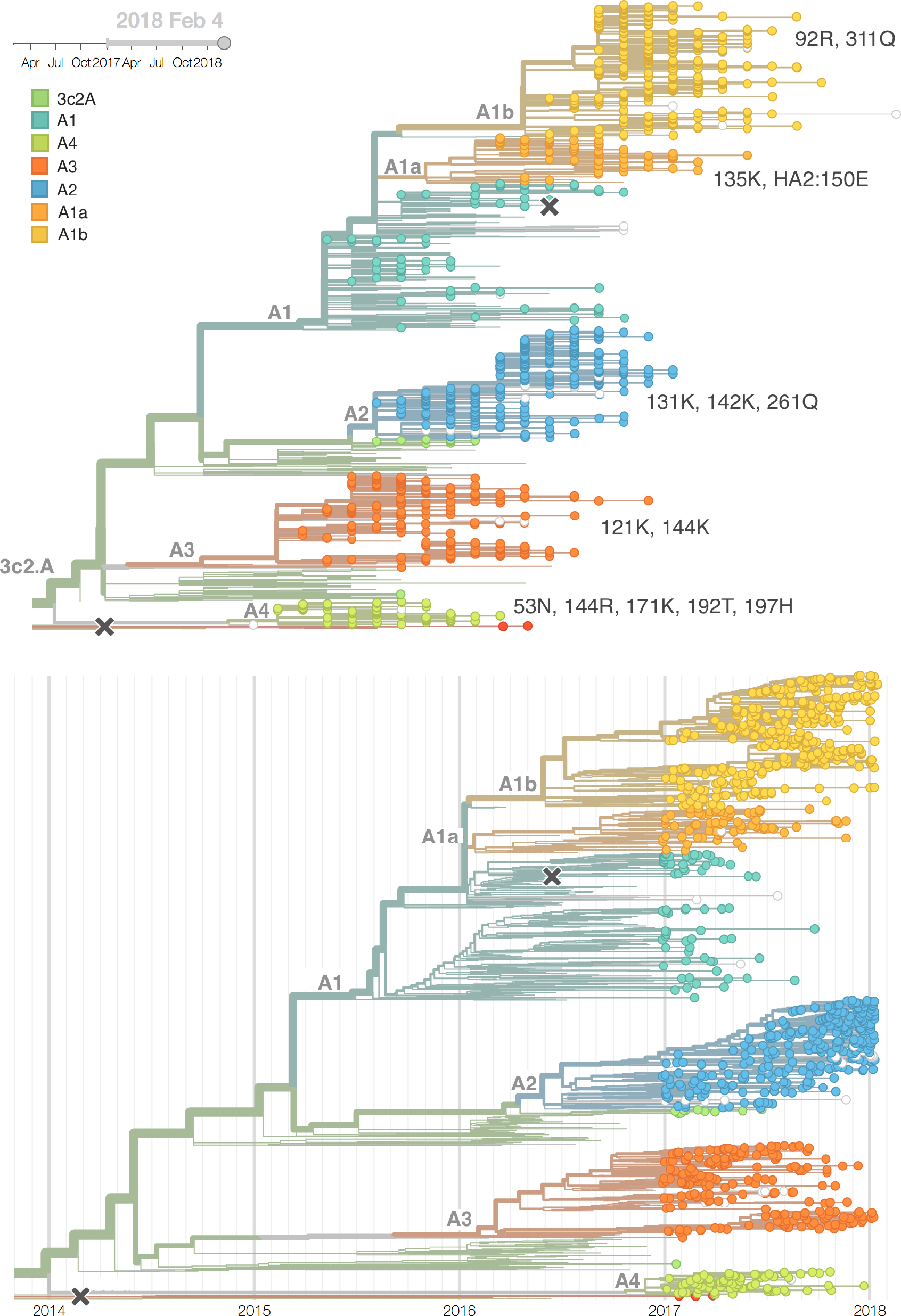
H3N2 / 3c2.A phylogeny colored by clade. We number these clades A1-A4. Each of these clades is non-nested and so all compete against each other. Also shown is a time calibrated phylogeny with equivalent coloring.

In the course of 2017, all of these clades had risen in frequency to dominate the H3N2 population (Fig. 3). However, clades A1a and A4 have recently declined leaving clades A1b, A2 and A3 to make up the majority of circulating viruses. Clade A4 was at high frequency in Oceania in the recent in Southern Hemisphere season, but is hardly observed in the current Northern Hemisphere season. Clade A1a is close to extinction as well. The remaining three clades (A1b, A2 and A3) have distinct geographic distributions. Clade A1b is estimated at ~30% of the global population, down from a peak of nearly 60%. Clade A2 is estimated at ~40% of the global population, but is dominating the current North American season and is increasingly observed in Europe. Clade A3 is estimated at a steady global frequency of about 25%.

**Figure 3.**
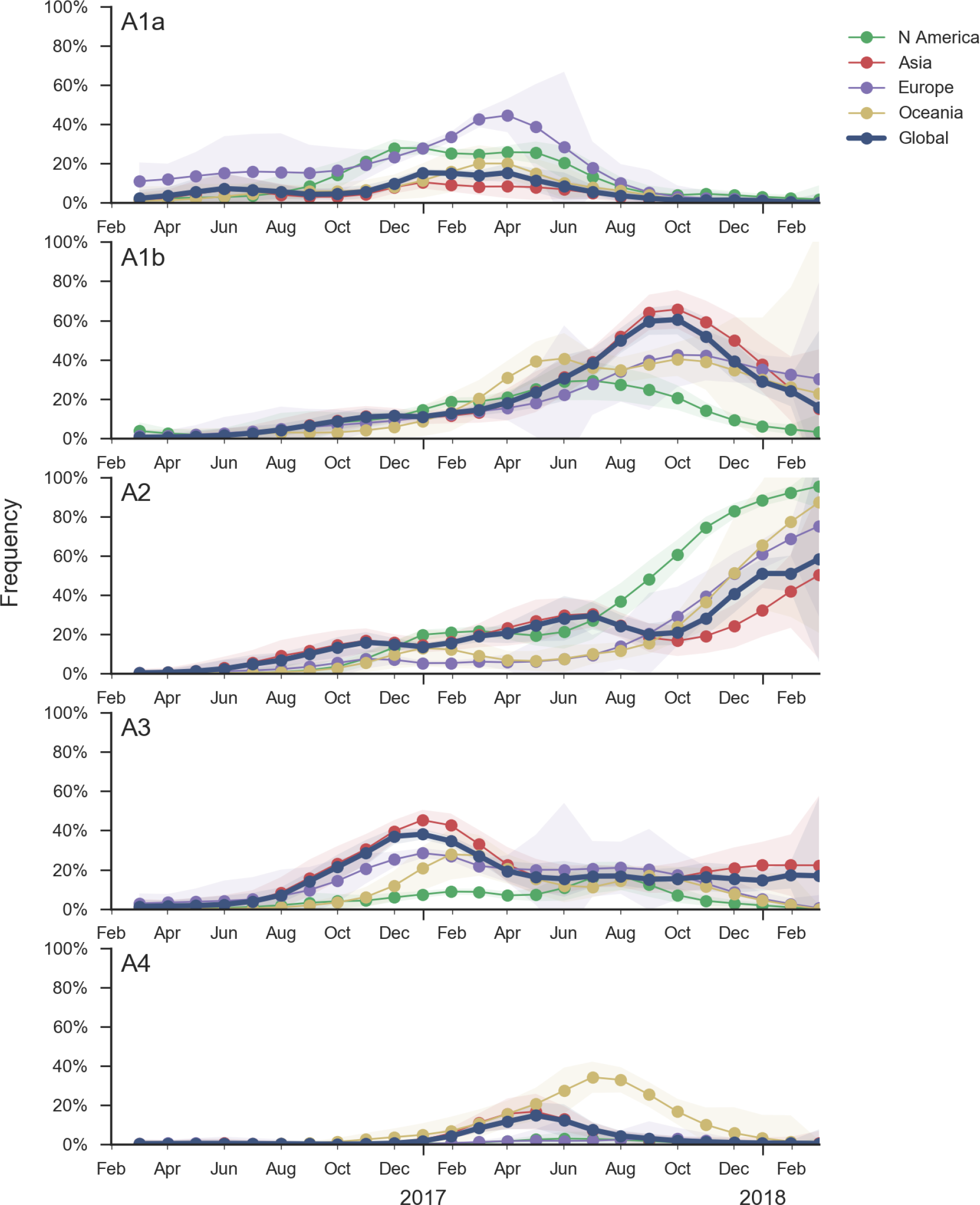
Frequency trajectories of 3c2.A subclades partitioned by clade then by region. We estimate frequencies of different clades based on sample counts and collection dates. Each clade is estimated according to a characteristic mutation unique to this clade. These estimates are based on all available data and global frequencies are weighted according to regional population size and relative sampling intensity. We use a Brownian motion process prior to smooth frequencies from month-to-month. Transparent bands show an estimate the 95% confidence interval based on sample counts.

Within these 5 clades, there have been frequent parallel mutations, particularly at HA1 sites 121, 135 and 142. The mutation T135K mutation in clade A3 is responsible for its recent resurgence in frequency. Clade A1b shows a similar pattern with mutations T135K and T135N making up two increasing subclades within the A1b clade.

Recent clade A2 viruses descend from a reassortment event about one year ago in which the HA and PB1 segments of a clade A2 virus moved into the background of a clade A1a virus (hence these viruses show up as HA clade A2) (Fig. 4). The neuraminidase of clade A1a differs from clade A2 by I176M, N329S, D339N, and P386S. The original neuraminidase of clade A2 and clade A3 viruses were outcompeted by the reassortant.

**Figure 4.**
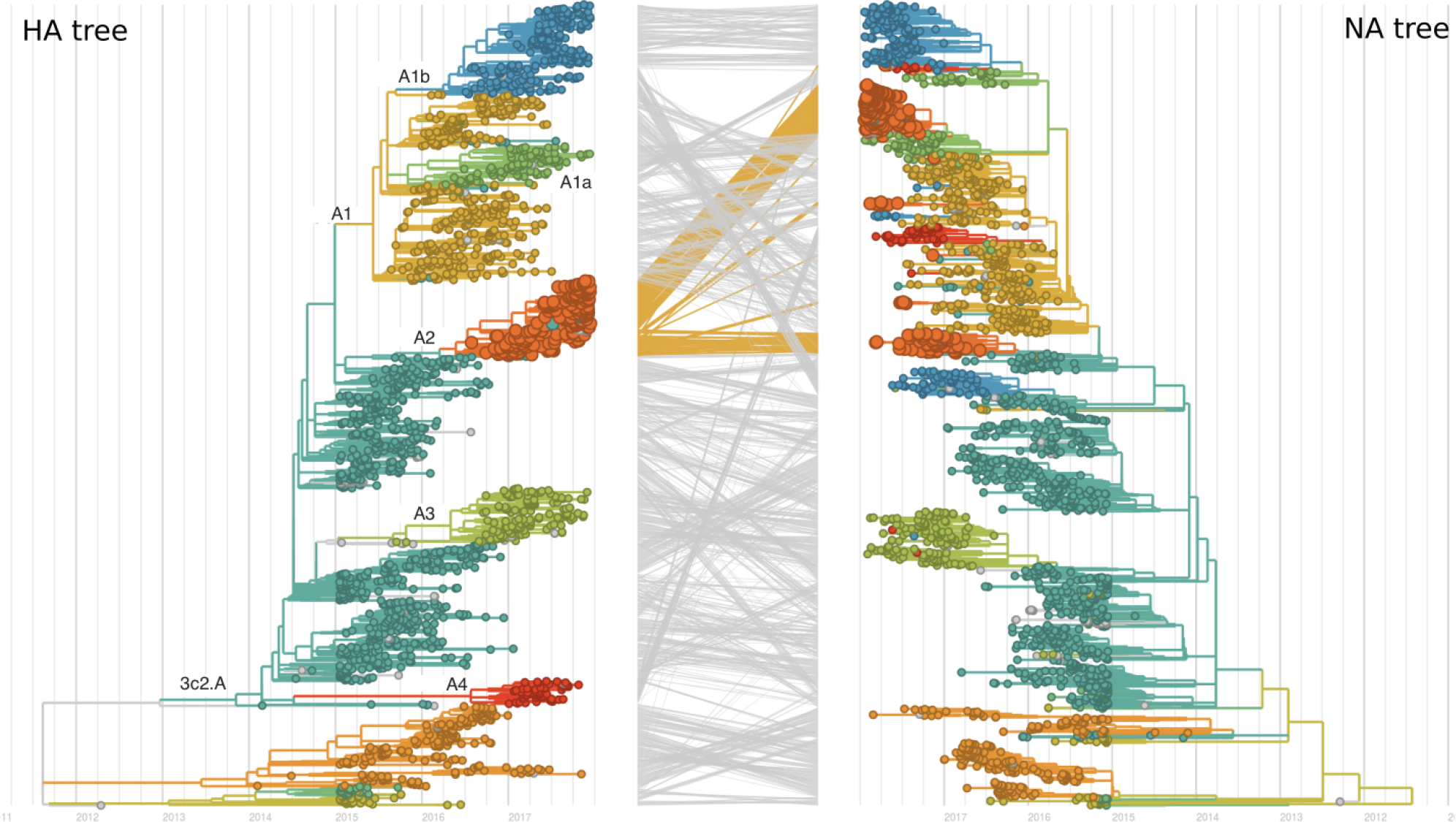
Reassortment patterns between HA and NA. Recent viruses from HA clade A2 are reassortants with clade A1a viruses. HA and PB1 derive from former clade A2, all other segments are derived from a clade A1a virus. 80% of recently circulating viruses carry an NA with N329S and several other substitutions. The only reference virus carrying this NA variant is A/Wisconsin/327/2017.

These four recently emerged variants constitute the most successful recent subclades (Fig. 5). Clade A1b viruses with 135K had increased rapidly in frequency in fall 2017, but are now declining globally. However, reassortant clade A2 viruses have completely driven the recent growth of A2 and these viruses are rapidly increasing in multiple regions.

**Figure 5.**
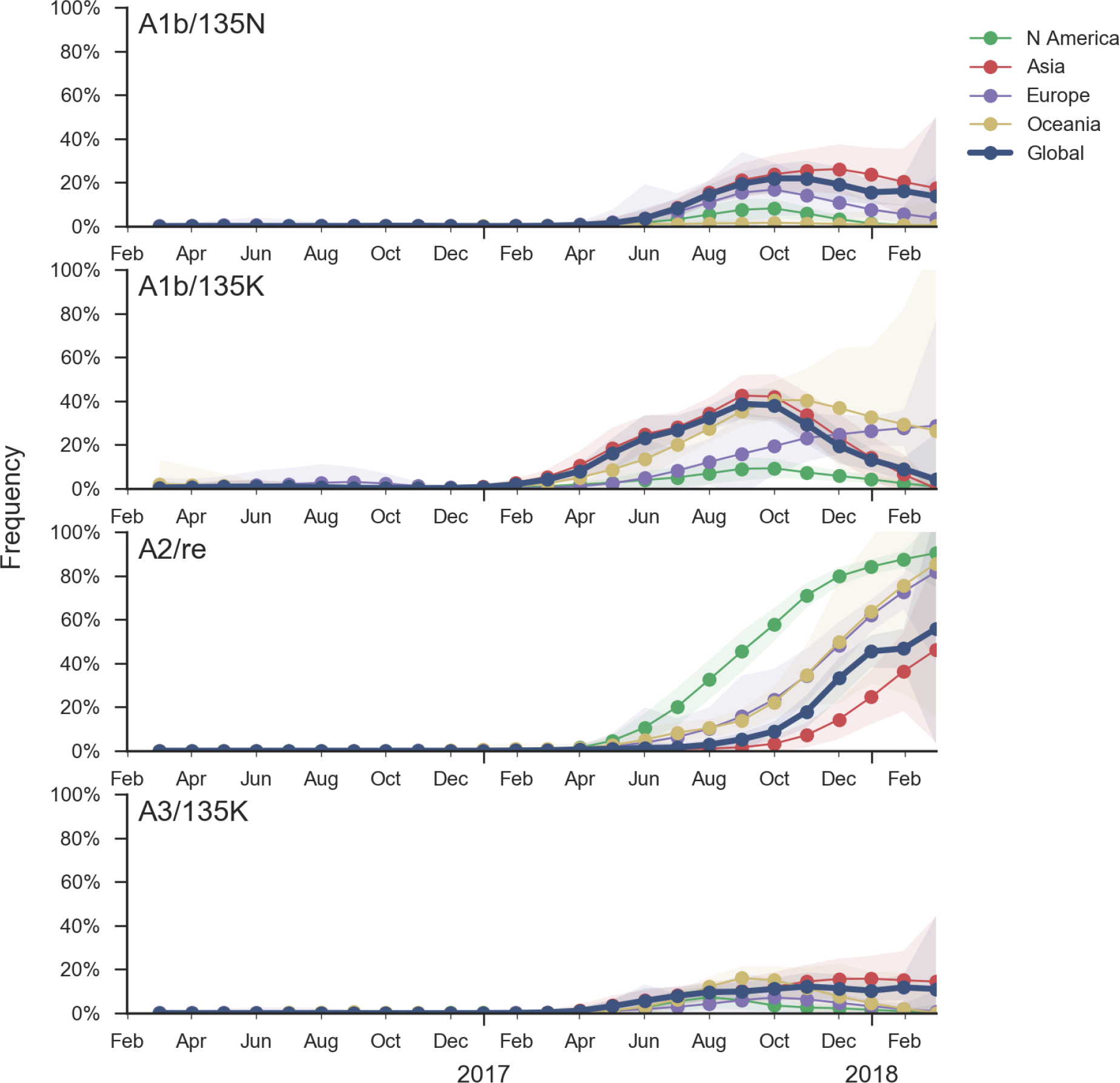
Frequency trajectories of emerging clades partitioned by clade then by region. We estimate frequencies of different clades based on sample counts and collection dates. Each clade is estimated according to a characteristic mutation unique to this clade. These estimates are based on all available data and global frequencies are weighted according to regional population size and relative sampling intensity. We use a Brownian motion process prior to smooth frequencies from month-to-month. Transparent bands show an estimate the 95% confidence interval based on sample counts.

## Antigenic properties

All 5 of these clades appear somewhat drifted according to antigenic analysis [2] of HI measurements provided by Influenza Division at the US Centers for Disease Control and Prevention (Fig. 6). Antigenic differences from the future Southern Hemisphere vaccine strain A/Singapore/Infimh-16-0019/2016 to circulating viruses are estimated to be between 0.4 and 2.2 antigenic units (or approximately a four-fold HI dilution).

**Figure 6.**
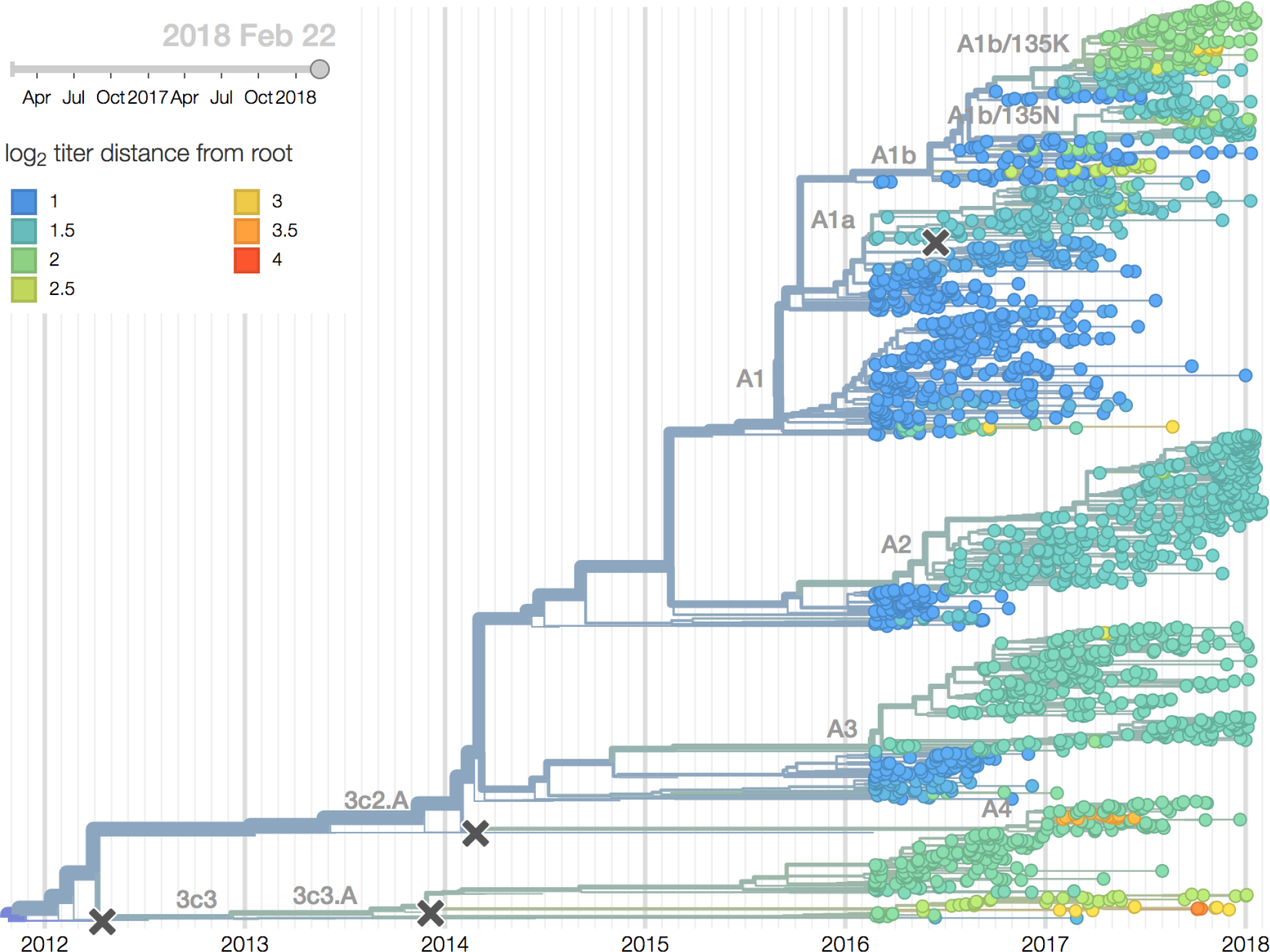
H3N2 phylogeny colored by antigenic distance to phylogeny root. Cooler color indicates greater antigenic similarity (less titer drop going from homologous to heterologous titers). More novel strains are greener and less novel strains are bluer. Estimates derive from HI data collected by the US Centers for Disease Control and Prevention.

## Projections

Recent behavior of cocirculating clades A1b, A2 and A3 would suggest that clades A1b and A2 are the most competitive. The highest local branching index (LBI) [3] is observed at the base in clade A1b concomitant with the T135K mutation (potential loss of glycosylation, minor antigenic impact) and in clade A2 following reassortment with an evolved NA background (Fig. 7). With clade A2 HA, there are no recent mutations that suggest adaptation, but the reassortment event has potentially changed intrinsic fitness and antigenic properties of NA. Clade A2 has risen extremely rapidly in North America, but it is not yet clear whether similar dynamics play out in Asia. However, while there is not large amounts of data, reassortant clade A2 viruses now appear to be spreading in Asia (Fig. 5). Clade A1b currently appears at higher frequency than clade A2 in Asia. However, clade A1b is declining, while clade A2 is increasing. The complete dominance of A2 viruses in North America may not strongly reflect future trends, as successful lineages often lead in Asia [4]. In a recent example, in spring 2016, 3c3.A viruses dominated towards the end of the North American season. However, these viruses did not successfully spread outside of North America.

**Figure 7.**
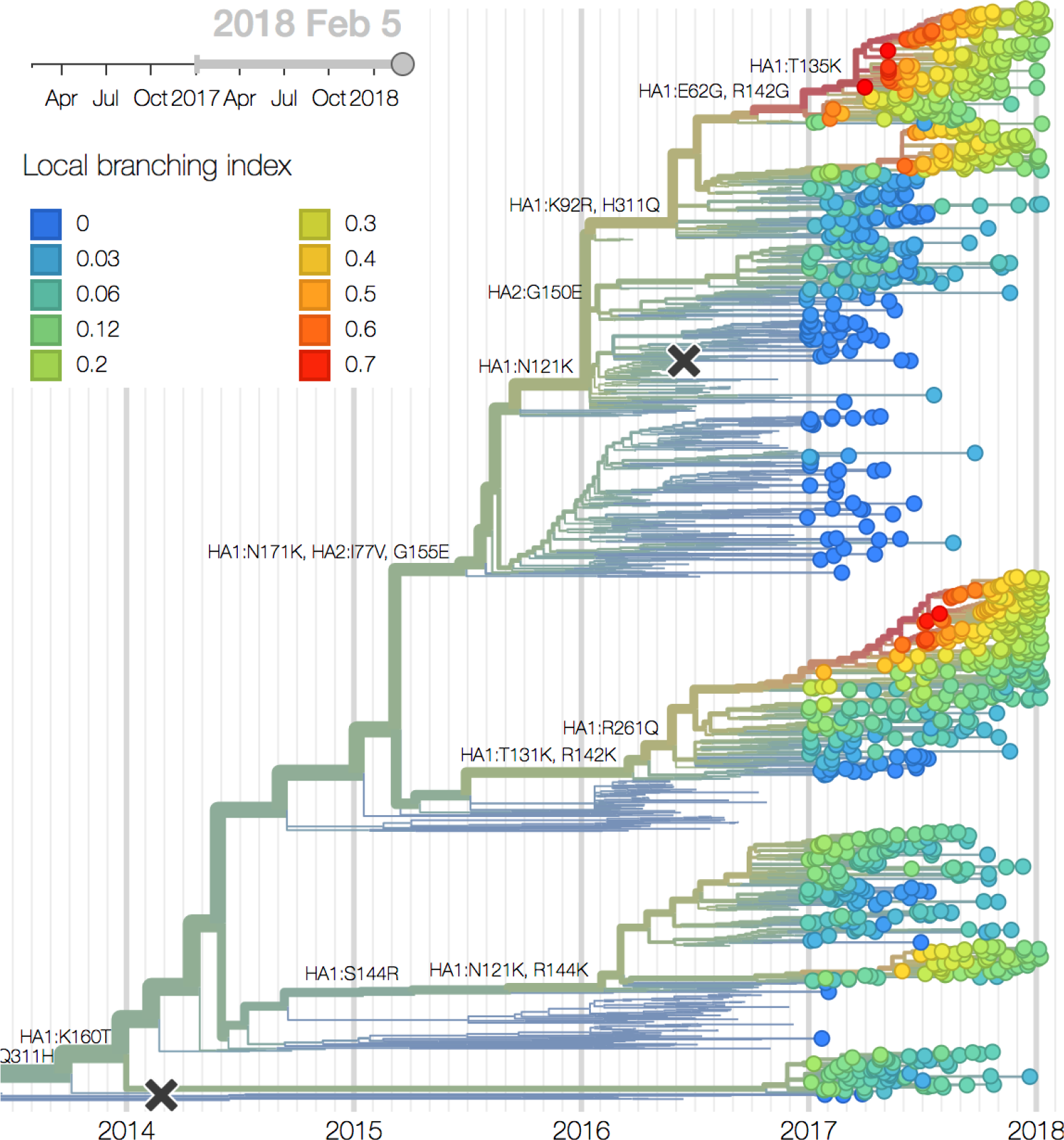
H3N2 / 3c2.A phylogeny colored by local branching index. The tree, zoomed into clade 3c2.A, is colored by local branching index. Redder nodes have seen faster recent growth. The branch with the T135K substitution in clade A1b and the branch leading up to the reassortment event in clade A2 have almost identical LBI.

**Figure 8.**
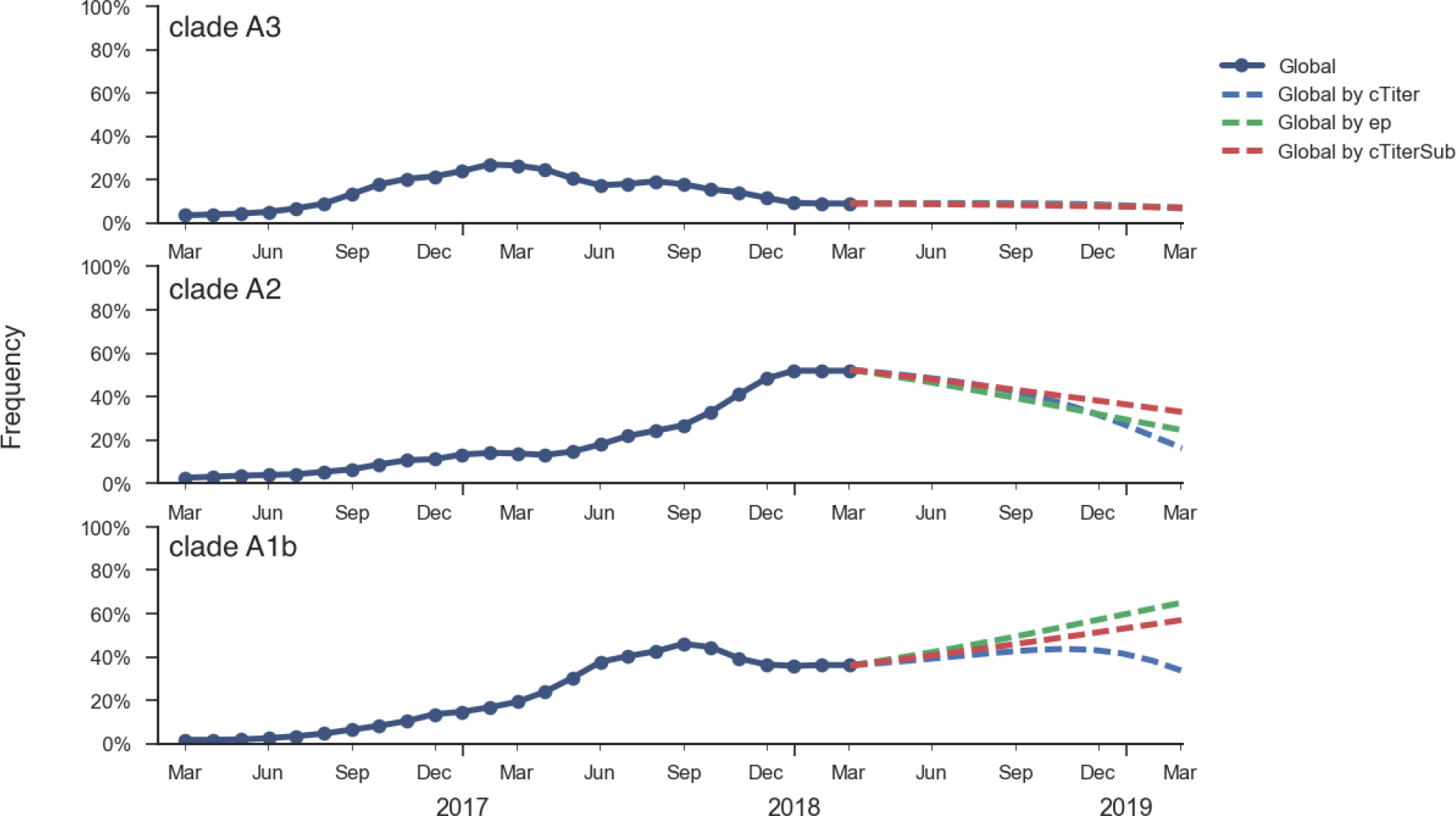
Forecast trajectories of clades A1b, A2 and A3. We compare 3 separate fitness predictors (epitope mutations ep, antigenic advancement based on the tree model `cTiter` and antigenic advancement based on the substitution model `cTiterSub`). Model parameters are tuned based fitting to previous 15 years of data.

Fitness correlates of epitope mutations [5] and antigenic phenotype estimated from HI data [2] suggest that intrinsic fitness of clade A1b viruses may be slightly greater than clade A2 viruses. Using these correlates in a formal fitness model, as originally described by Łuksza and Lässig [6], suggests that clade A1b viruses will increase in frequency over the coming year and clade A2 viruses will decrease. However, fitness differences between these two clades are small and the model predicts that both will still circulate in Feb 2019. Additionally, this fitness model cannot capture the effects of the reassortment event and so it may not be properly weighting clade A2 viruses.

We think that clade A1b viruses or reassortant clade A2 viruses are most likely to increase in the coming year. It is difficult to project a winner and cocirculation is a likely outcome.

## A/H1N1pdm

A clade comprising mutations S74R, S164T and I295V has recently swept to fixation. The rapidity of this sweep suggests a selective origin. However, there is no evidence of antigenic change.

We base our primary analysis on a set of viruses collected between Mar 2016 and Jan 2018, comprising approximately 100 viruses per month during relevant months of 2017 (Fig. 9). We use all available data when estimating frequencies of mutations and weight samples appropriately by regional population size and relative sampling intensity to arrive at a putatively unbiased global frequency estimate. Phylogenetic analyses are based on a representative sample of about 2000 viruses.

**Figure 9.**
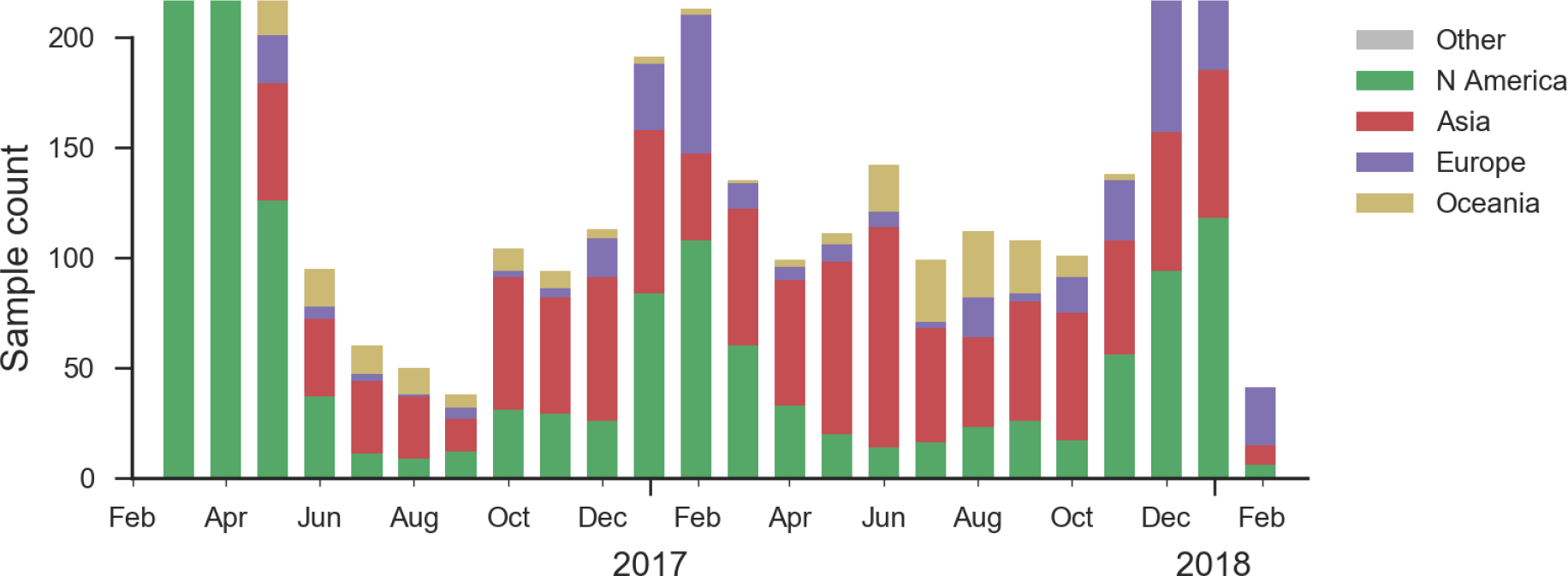
A/H1N1pdm sample counts through time and across regions. This is a stacked bar plot, so that in many months there are more than 100 samples.

H1N1pdm recently underwent a selective sweep of the 6b.1 clade comprising mutations S162N and I216T. All currently circulating H1N1pdm viruses are clade 6b.1. The H1N1pdm vaccine strain was updated in Sep 2016 to A/Michigan/45/2015 to match 6b.1 viruses.

Within 6b.1 there has emerged a subclade comprising S74R, S164T and I295V that has recently swept to fixation (Fig. 10, Fig. 11). Subsequently, several amino acid substitutions have emerged within the S164T clade. The S183P substitution was circulating outside of the S164T clade previously and now reemerged within the S164T clade, predominantly in North America. Furthermore, the T120A mutations has risen to about 40% frequency over the last few months.

**Figure 10.**
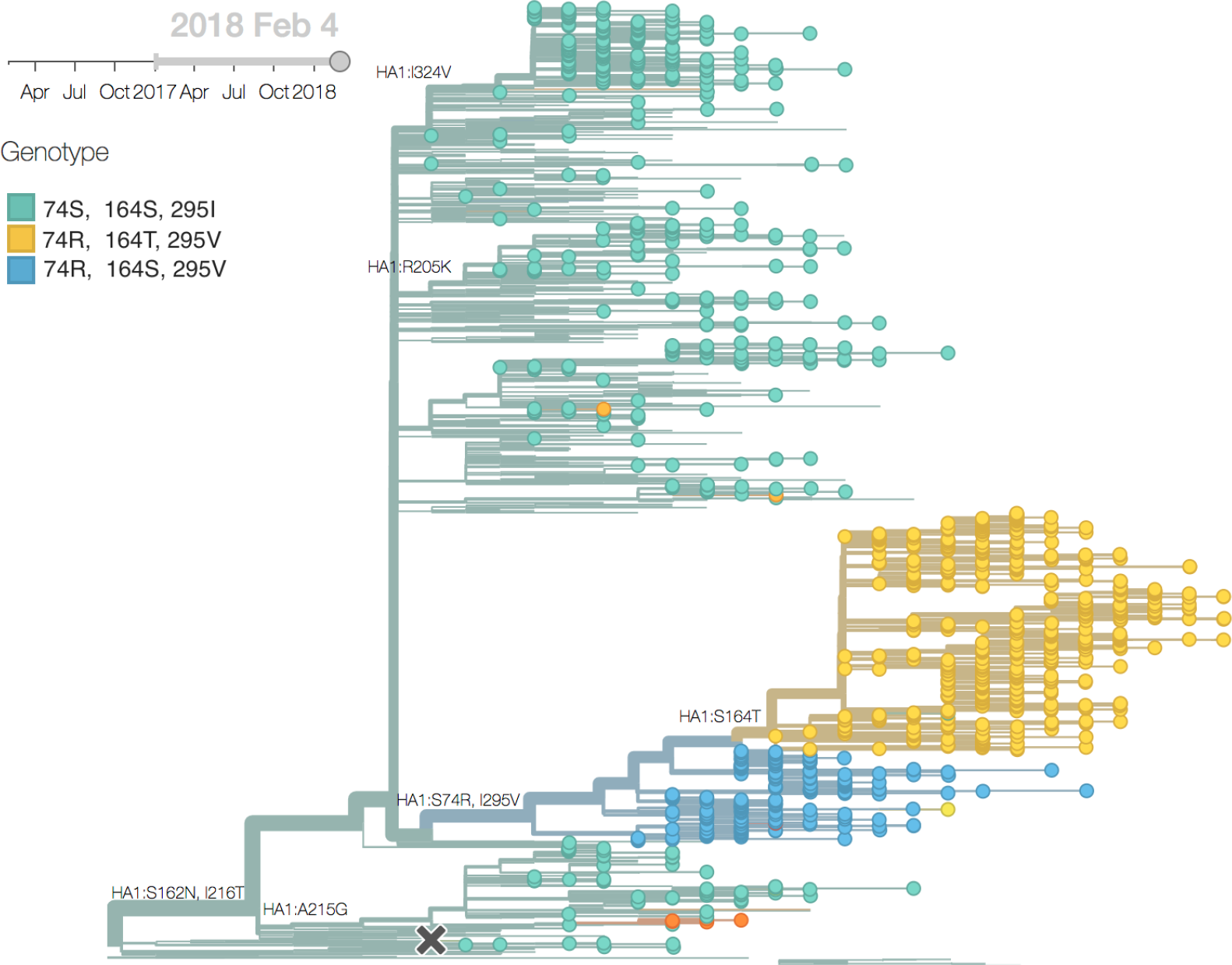
H1N1pdm phylogeny colored by genotype.

**Figure 11.**
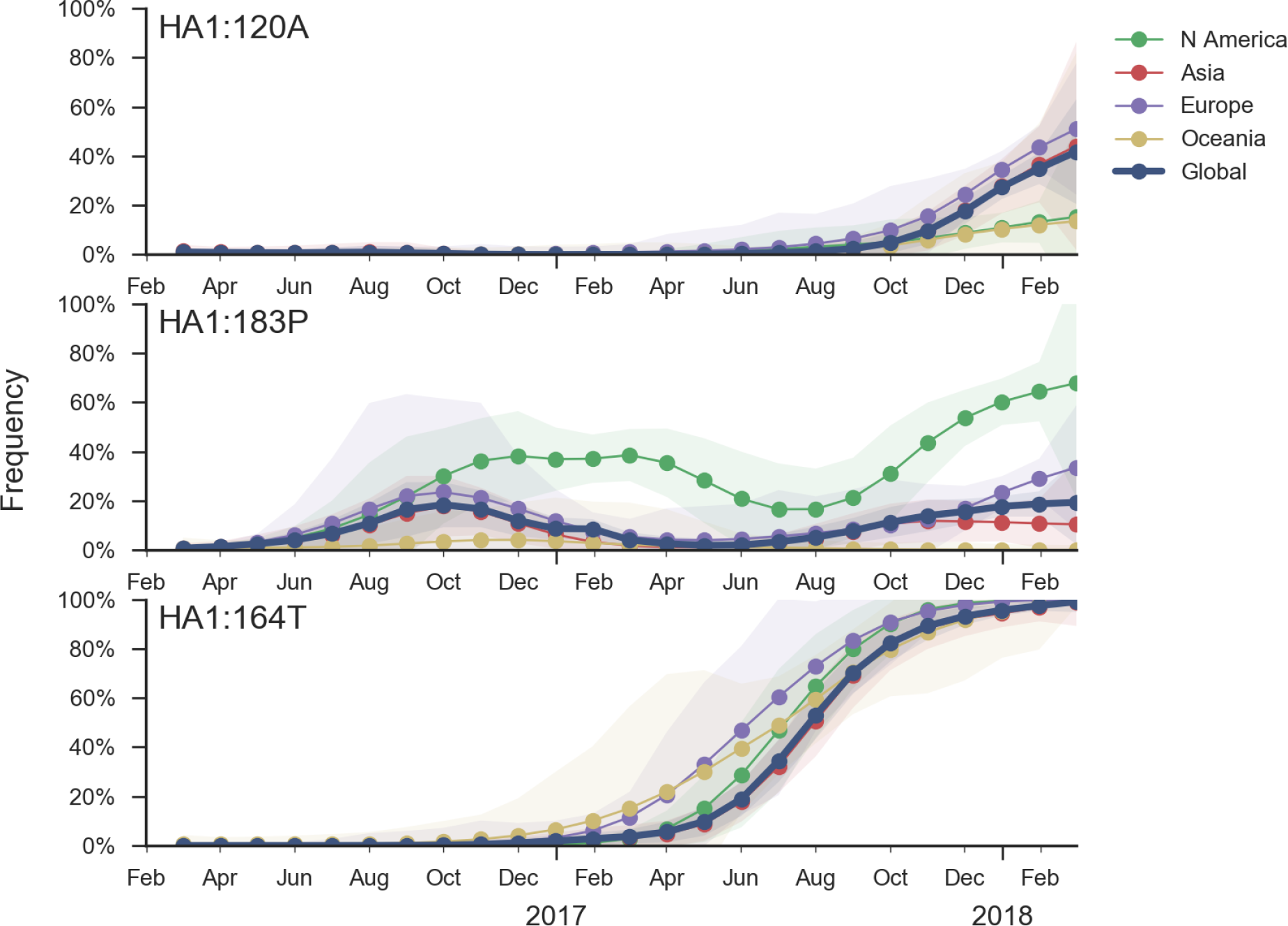
Frequency trajectories of recent mutations in A/H1N1pdm viruses. We estimate frequencies of different mutations based on sample counts and collection dates. These estimates are based on all available data and global frequencies are weighted according to regional population size and relative sampling intensity. We use a Brownian motion process prior to smooth frequencies from month-to-month. Transparent bands show an estimate the 95% confidence interval based on sample counts.

In a pattern familiar in the recent history of H1N1pdm evolution, none of these mutations has obvious an obvious impact in antigenic essays (Fig. 12), yet they exhibit rapid shifts in frequency and recurrent mutations are successful on different genetic backgrounds, suggesting potential selective origins.

**Figure 12.**
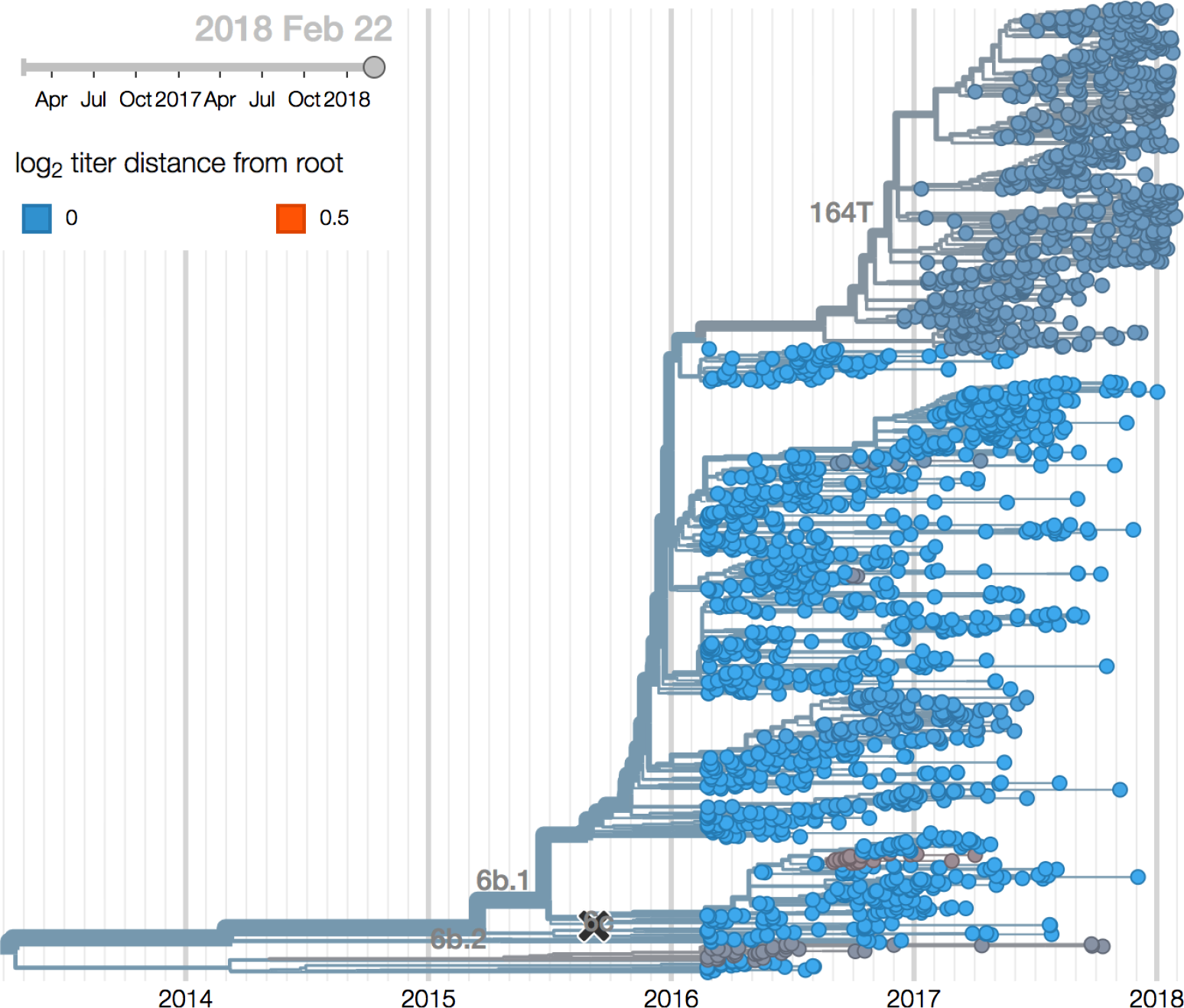
H3N2 phylogeny colored by antigenic distance to phylogeny root. Cooler color indicates greater antigenic similarity (less titer drop going from homologous to heterologous titers). More novel strains are greener and less novel strains are bluer. Estimates derive from HI data collected by the US Centers for Disease Control and Prevention.

## B/Vic

Very little B/Vic activity has been observed in recent months. A clade with a two codon deletion at sites HA1:162/163 has gradually risen in frequency. HI measurements suggest an 8 to 16-fold titer drop relative to the vaccine strain, but this antigenic change has not yet resulted in a rapid rise of this variant.

We base our primary analysis on a set of viruses collected between Mar 2016 and Jan 2018, comprising >150 viruses per month in Dec 2016 to Apr 2017 and 20-50 viruses per month more recently (Fig. 13). We use all available data when estimating frequencies of mutations and weight samples appropriately by regional population size and relative sampling intensity to arrive at a putatively unbiased global frequency estimate. Phylogenetic analyses are based on a representative sample of about 2000 viruses.

**Figure 13.**
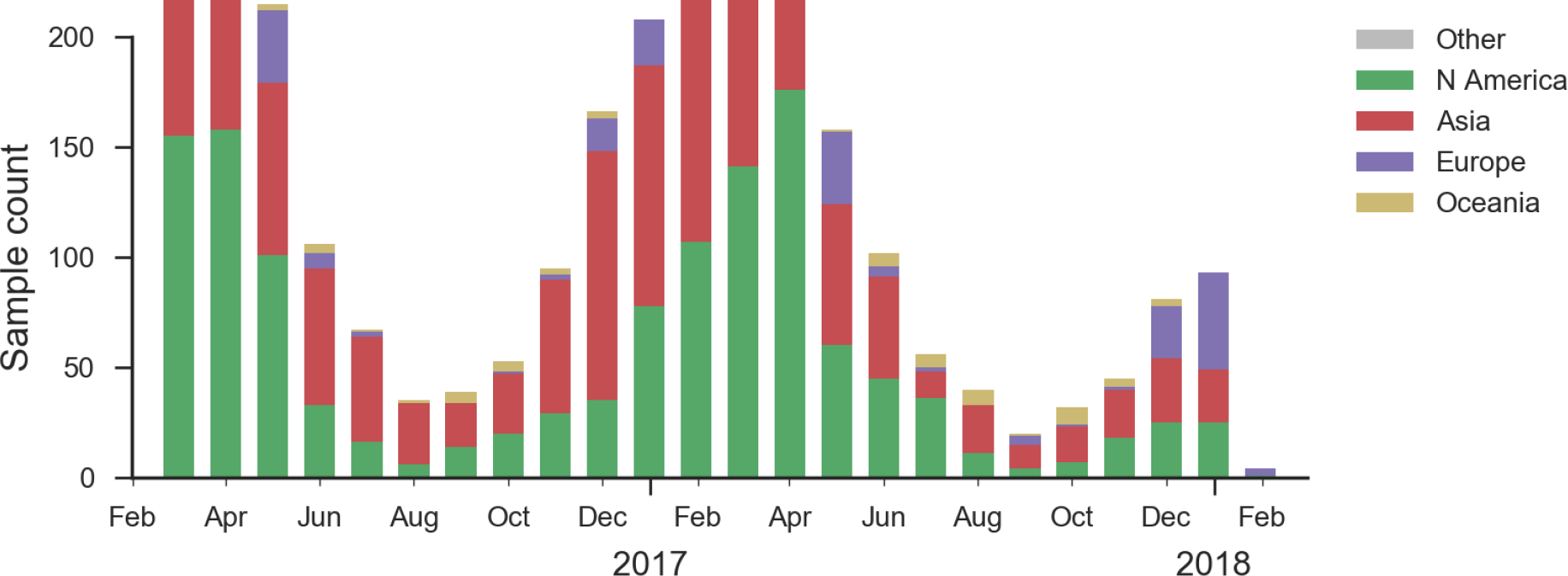
B/Vic sample counts through time and across regions. This is a stacked bar plot, so that in many months there are more than 200 samples.

Recent B/Vic sequences fall into three noteworthy clades. Most importantly, a clade defined by a deletion of codons 162/163 along with mutations D129G and I180V emerged in 2016 and has been slowly spreading since (Fig. 14). It is now at ~17% global frequency and appears at high frequency in recent viruses from Europe and North America, although these estimates are based on little recent data (Fig. 15). Additionally, a clade defined by a I175V substitution in HA1 has slowly risen to moderate frequency in North America. Another clade defined by mutation K209N has been circulating at low frequency in Asia throughout last year.

**Figure 14.**
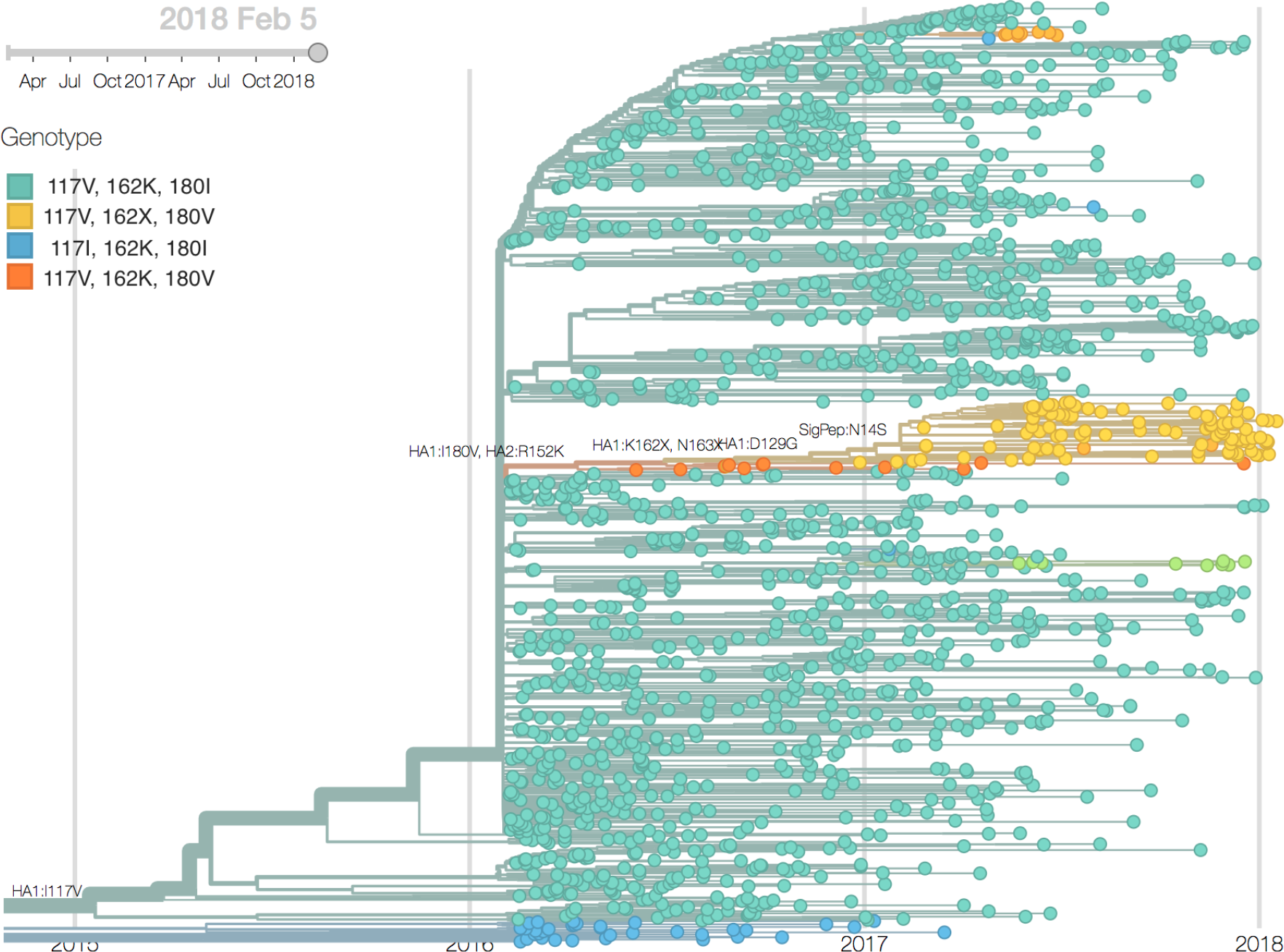
B/Vic phylogeny colored by genotype at sites HA1 117, 162 and 180. The major circulating variant within B/Vic is the 162X, 163X deletion variant.

**Figure 15.**
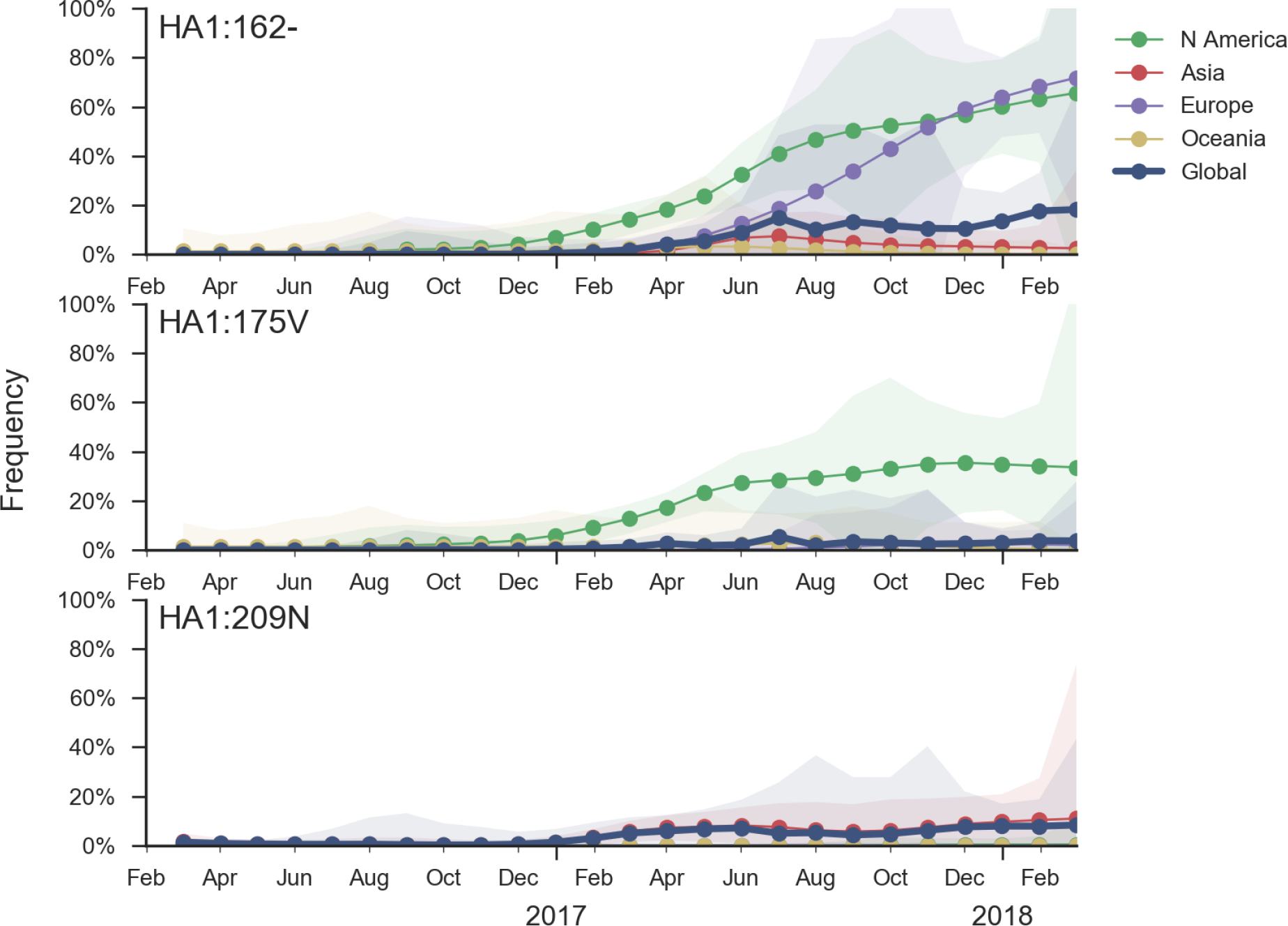
Frequency trajectories of recent mutations in B/Vic viruses. We estimate frequencies of different mutations based on sample counts and collection dates. These estimates are based on all available data and global frequencies are weighted according to regional population size and relative sampling intensity. We use a Brownian motion process prior to smooth frequencies from month-to-month. Transparent bands show an estimate the 95% confidence interval based on sample counts. All estimates are fairly noisy, as few sequences are available.

The deletion variant (DV) shows a strong antigenic impact with an 8 to 16-fold titer drop relative to B/Texas/2/2013 (Fig. 16). This would suggest increased fitness, however, the slow spread of the variant suggests that it may have instrinsic deficiencies in viral fitness. Still, if current trends continue, then the deletion variant is expected to slowly increase in frequency over the coming year. An extremely simple logistic growth model where growth rate is calculated from increase in frequency over the past 12 months, places frequency of the deletion variant at ~50% in Jan 2019 (Fig. 17). We caution again over-interpretation of this model; our expectation is that the deletion variant will circulate at moderate frequency in the coming year. Overall, there was very little B/Vic activity resulting in few sequenced viruses; this reduces confidence in these patterns.

**Figure 16.**
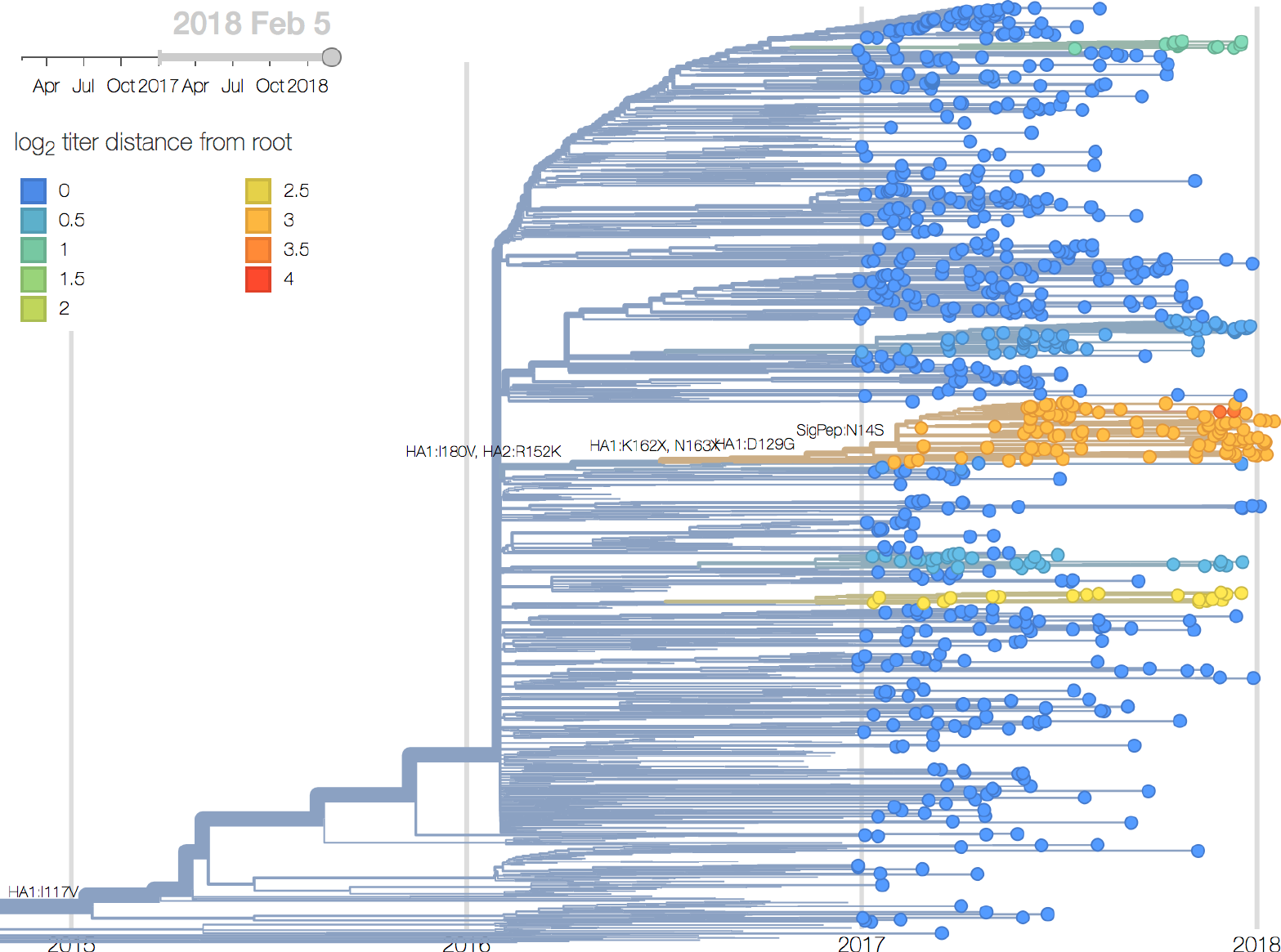
Vic phylogeny colored by antigenic distance. This shows estimates of HI titer drop to root of the phylogeny from titer model. Cooler color indicates greater antigenic similarity (less titer drop going from homologous to heterologous titers).

**Figure 17.**
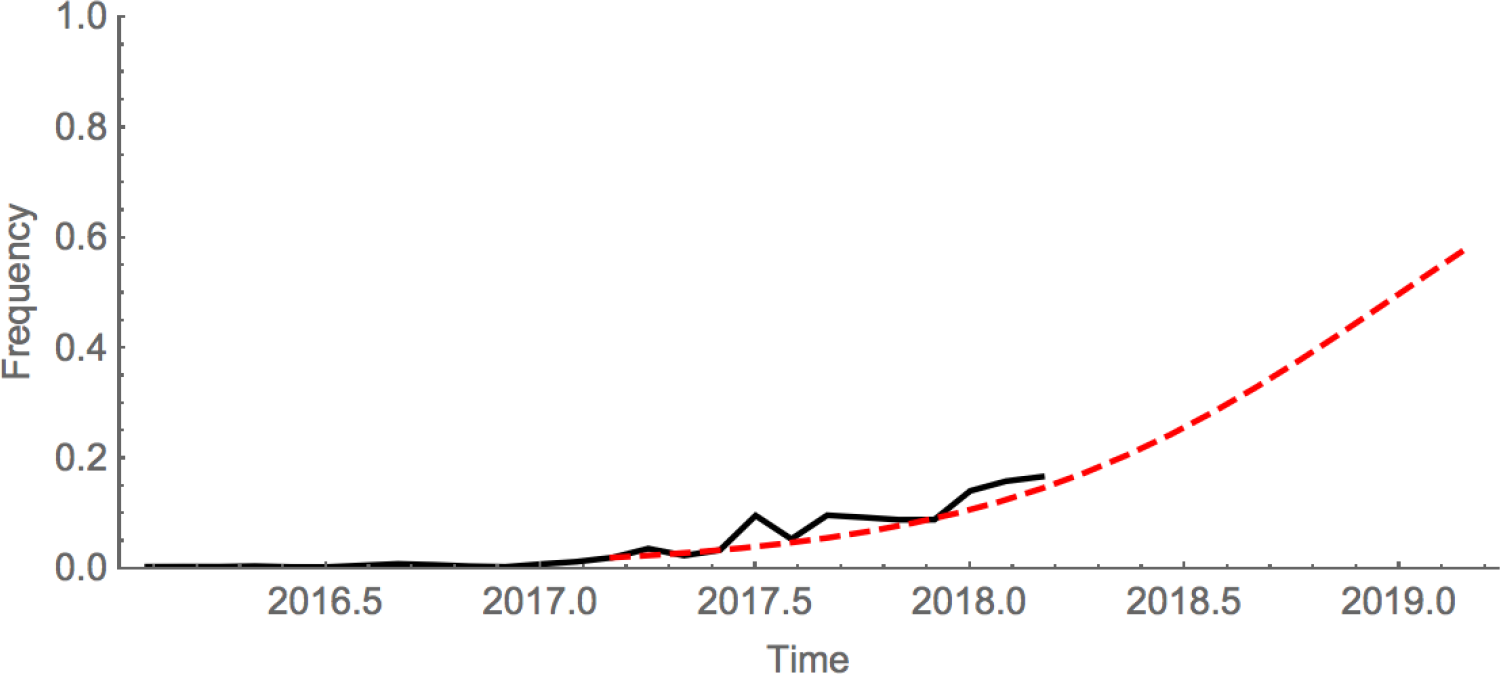
Logistic growth model of spread of deletion variant. Growth rate is estimated from increase in the past 12 months and future growth a simple extrapolation.

## B/Yam

Europe experienced a strong and early B/Yam season in absence of amino acid variation in HA or antigenic diversity. However, several mutations in NA have rapidly swept or risen to intermediate frequencies.

We base our primary analysis on a set of viruses collected between Mar 2016 and Jan 2018, comprising between 50 and 150 viruses per month during recent months of 2017 and 2018 (Fig. 18). The majority of recent B/Yam sequences are from Europe. We use all available data when estimating frequencies of mutations and weight samples appropriately by regional population size and relative sampling intensity to arrive at a putatively unbiased global frequency estimate. Phylogenetic analyses are based on a representative sample of about 2000 viruses.

**Figure 18.**
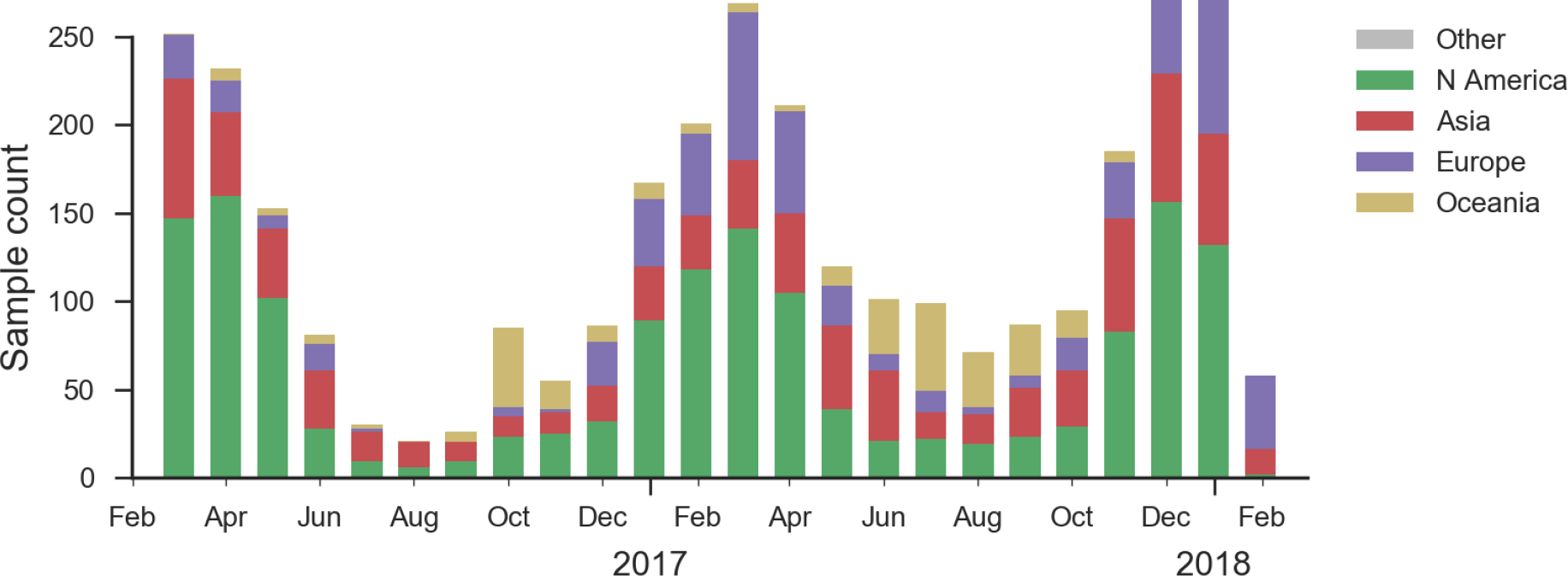
Sample counts through time and across regions. This is a stacked bar plot, so that in many months there are ~200 total samples.

We observe very little variation among the HA segments of B/Yam viruses (Fig. 19). Previously circulating variants have either gone to fixation (M251V) or gone extinct (K211R). This lack of amino acid variation is consistent with little antigenic variation in HI assays.

**Figure 19.**
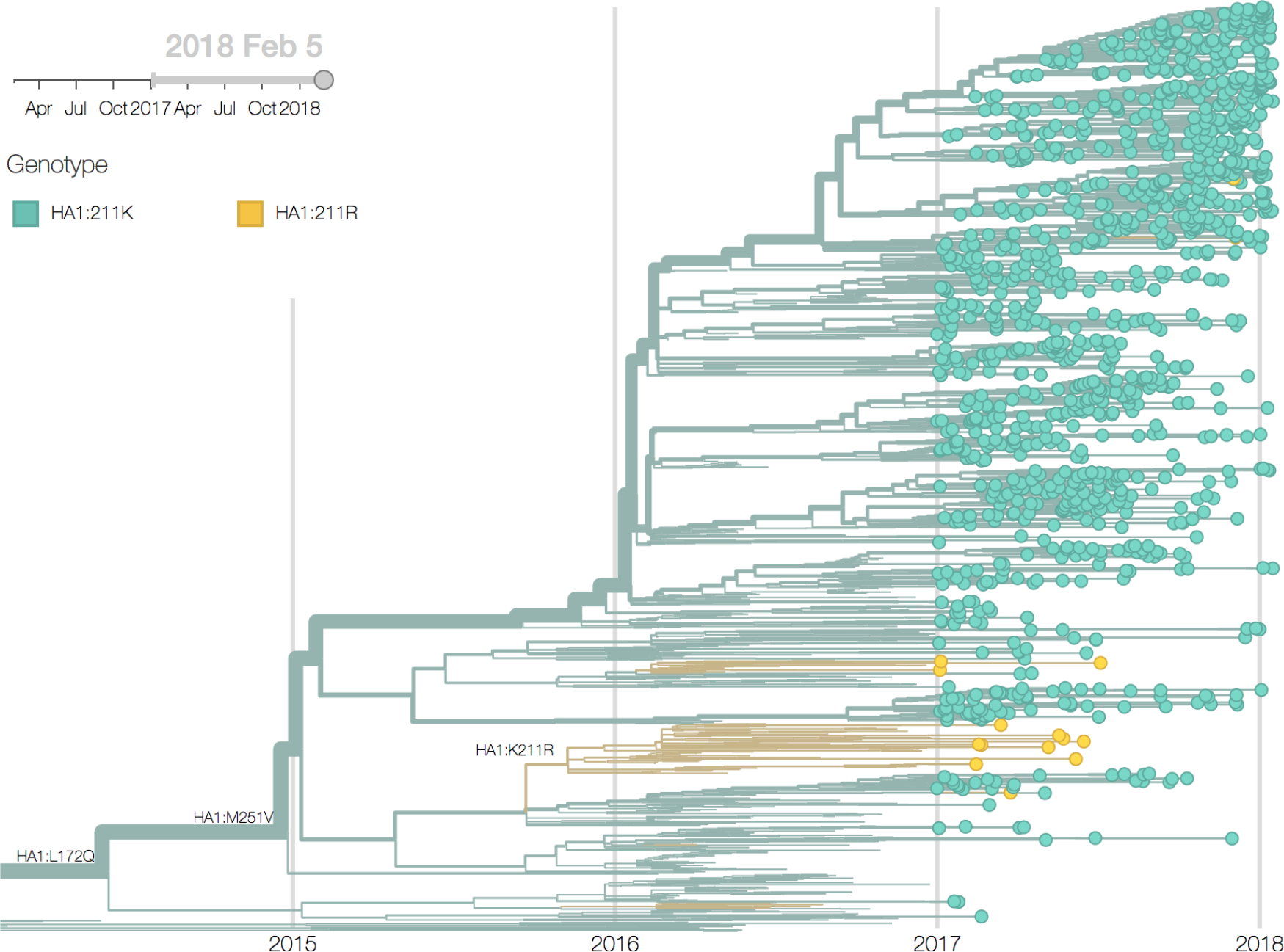
B/Yam phylogeny colored by genotype at site HA1 211. We observe very little diversity in Yam HA.

While little amino acid variation is observed in recent HA sequences, a number of substitutions in NA have risen to high frequency (Fig. 20). The mutation 342K rose from low frequency a year ago close to fixation. Within this clade, the mutation S402P is globally at about 50% frequency, dominating Europe. Outside of the clade defined by 402P, two mutations rose to 10-20% since the summer: S246T is common in Asia, while A395S is more common in North America (Fig. 21). At sites 395 and 342, we observe several independent mutations with significant circulation during the last 3 years. The significance of these mutations is unclear to us, but their rapid rise, recurrent mutation, and the unusual early dominance of B/Yam in Europe suggests a selective origin.

**Figure 20.**
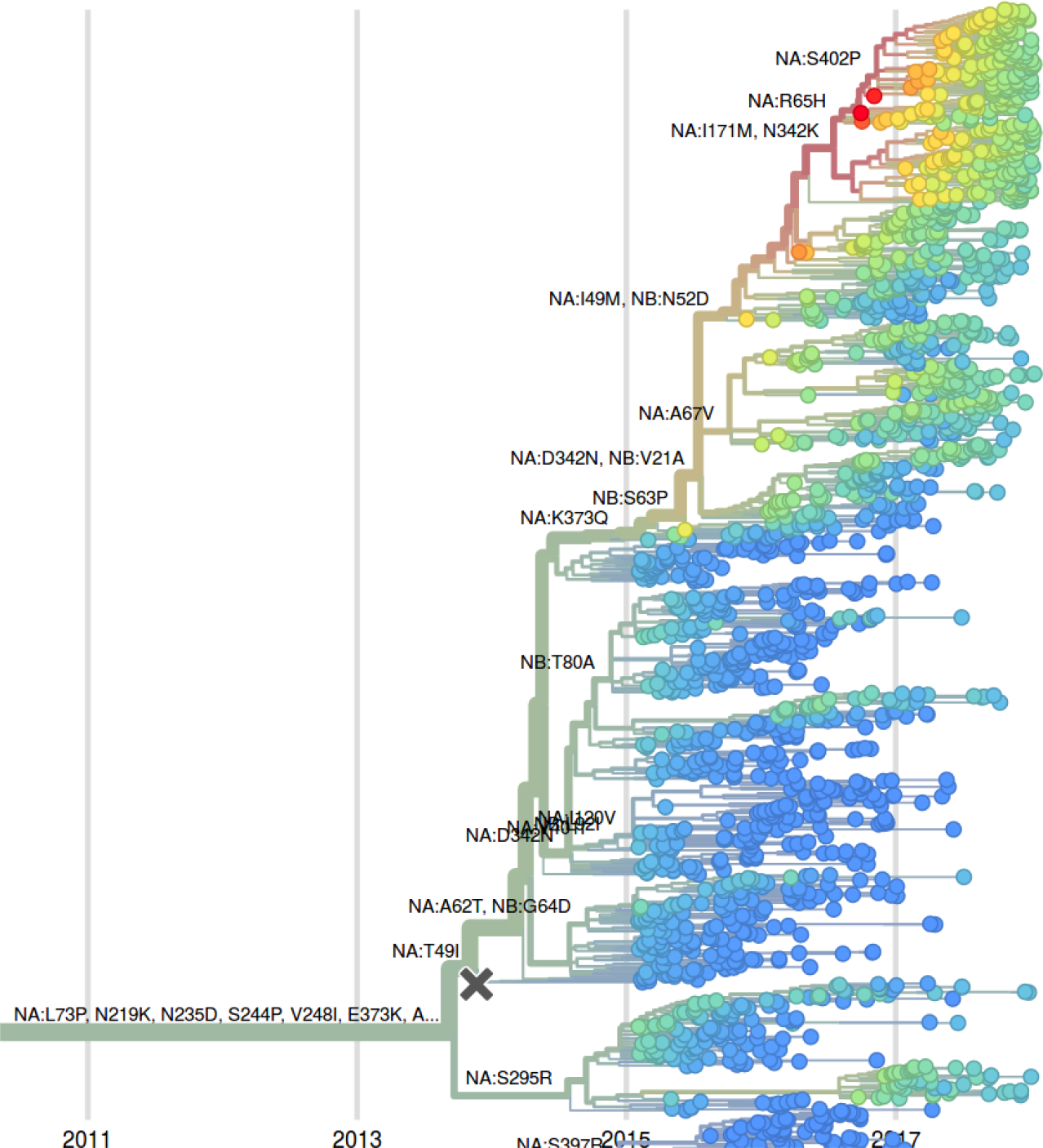
Phylogenetic tree of NA sequences of recent B/Yamagata isolates, colored by LBI. NA has undergone a number of recent amino acid substitutions, some of which have rapidly risen to high frequencies. With the exception of a small subclade at the bottom of the tree, almost all recent sequences fall into a clade carrying substitutions I171M and N342K.

**Figure 21.**
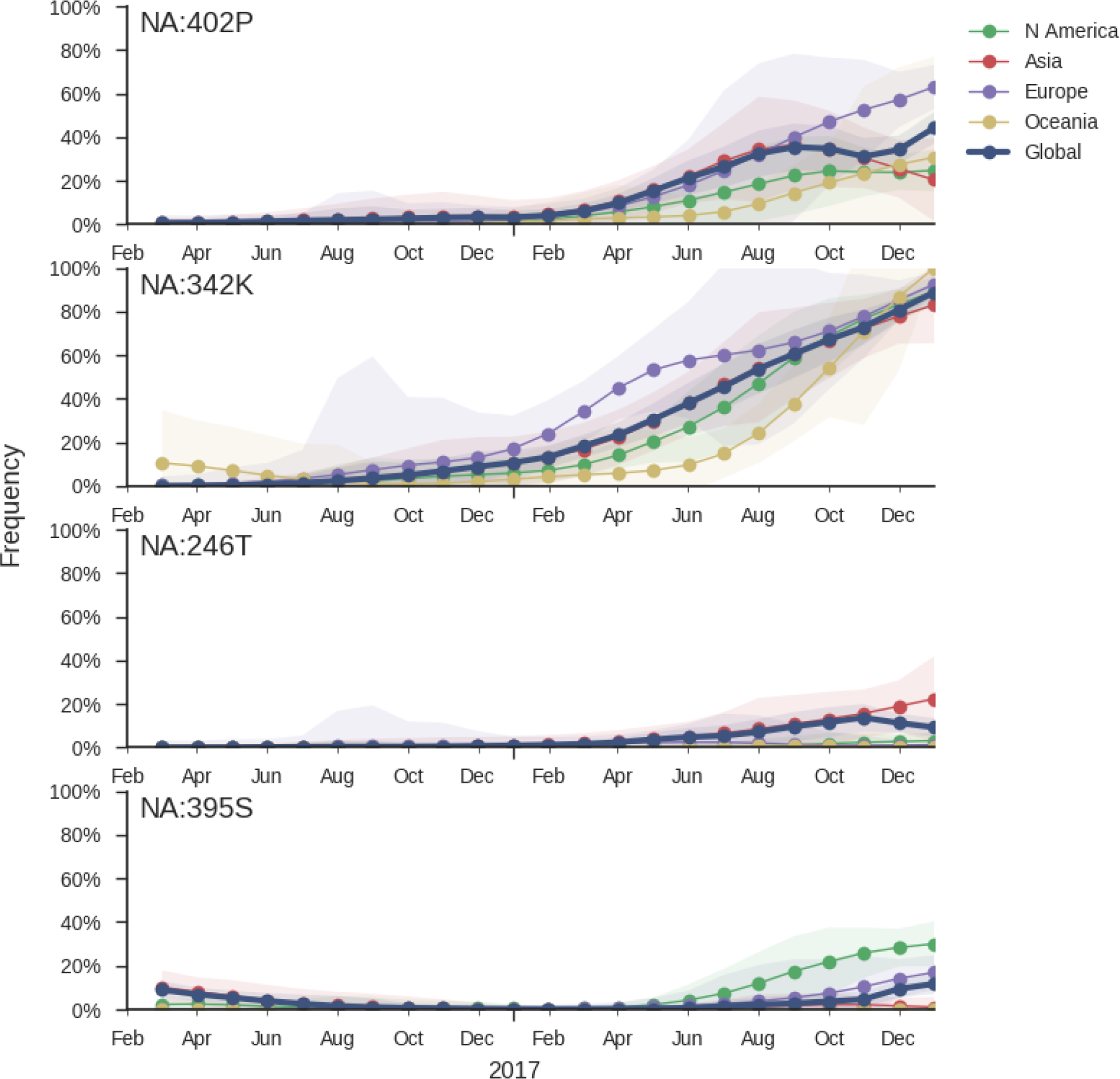
Frequency trajectories of substitution in NA of B/Yam. We estimate frequencies of different mutations based on sample counts and collection dates. The substitutions 402P, 246T, and 395S are sister clades. Transparent bands show an estimate the 95% confidence interval based on sample counts.

The LBI, when applied to the NA tree of B/Yam, suggests that the clade defined by the N342K mutation will take over (it is already above 80%) and that within this clade the subclade defined by the R65H will dominate. However, the dominance of isolates from European might skew this inference.

## Acknowledgements

We thank the Influenza Division at the US Centers for Disease Control and Prevention, the Victorian Infectious Diseases Reference Laboratory at the Australian Peter Doherty Institute for Infection and Immunity, the Influenza Virus Research Center at the Japan National Institute of Infectious Diseases, the Crick Worldwide Influenza Centre at the UK Francis Crick Institute for data sharing and feedback. We thank Barney Potter for data cleaning and organization and we thank John Huddleston for fitness model implementation. We thank David Wentworth, Rebecca Garten and Xiyan Xu for insight regarding analysis directions.

